# Advancing eDNA methods for monitoring the reproduction of quagga and zebra mussels in lakes

**DOI:** 10.1101/2025.09.25.678458

**Authors:** Marine Vautier, Isabelle Domaizon

## Abstract

This study demonstrates the applicability of environmental DNA (eDNA) methods to monitor the reproductive periods of two invasive freshwater mussels, *Dreissena polymorpha (*zebra mussel) and *Dreissena rostriformis bugensis (*quagga mussel), addressing the need for improved detection and monitoring techniques for these ecosystem-altering species.

New primers and probes for multiplex droplet digital PCR (ddPCR) were developed to enable the discrimination between zebra and quagga mussels, whose veliger larvae are morphologically indistinguishable. Three eDNA sampling methods were compared: integrated water samples (0-50 m depth), sub-surface water samples, and plankton bulk samples. These methods were applied in four deep peri-alpine lakes and the results obtained with eDNA were compared to traditional veliger microscopic counts.

The values obtained with the three eDNA approaches are positively corelated with visual counting of veliger larvae, but integrated water and plankton bulk eDNA samplings were found to be more effective in quantifying dreissenid veliger larvae and estimating reproductive periods than sub-surface water eDNA. The bulk-based approach is robust for qualitative presence/absence veliger larvae information, with a positive eDNA signal only when larvae are observed; it is the only eDNA method that successfully detected zebra signals in Lake Geneva. However, an overestimation of winter reproduction signal was observed for one lake with this method. The water-integrated approach captures well the quantitative dynamics of counted veliger larvae, but with the presence of false positives reproductive signal (mussel eDNA signal when no larvae are observed). The eDNA approaches tested here have enabled the characterisation of the reproductive dynamics of each of the two species in the studied lakes, highlighting the fact that quagga can reproduce throughout the year in peri-alpine lakes where it is dominant, whereas in lakes dominated by zebra, the reproduction period is limited to the warm season. The study also provides the first clue of quagga mussel presence in Lakes Annecy and Aiguebelette.

Although further methodological refinements are suggested for the eDNA approaches evaluated in this study, those approaches are nonetheless already applicable and can contribute to a better understanding of these invasive species ecology in lakes.

## Introduction

*Dreissena polymorpha* (zebra mussel) and *Dreissena rostriformis bugensis* (quagga mussel) are two freshwater mussel species native from the Ponto-Caspian region, recognized as invasive alien species (IAS) due to their global spread and ecological impacts (Son, 2007). These species are notorious invaders of freshwater ecosystems worldwide because of their rapid expansion, high reproductive rates, and detrimental effects on native biodiversity and ecosystem functioning (reviewed in Nalepa & Schloesser, 1992, 2014; Karatayev et al., 2002, 2015). Both zebra and quagga mussels are filter feeders capable of processing substantial volumes of water, extracting phytoplankton, suspended particles, and nutrients. Consequently, their presence significantly alters benthic habitats, affecting invertebrate communities, while also impacting planktonic communities, trophic relationships, and nutrient cycling (Karatayev et al., 2007; Budd et al., 2001; Raikow et al., 2004; Vanderploeg et al., 2010; Rowe et al., 2015; Beekey et al., 2004). While zebra mussels have been present in Western Europe since the 19th century, invading Alpine and sub-Alpine lakes in the 1960s (Pollux et al., 2010), the arrival of quagga mussels is more recent. Quagga first appeared in Western Europe from the mid-2000s (Bij de Vaate et al. 2013), with a theoretical appearance from 2014 in deep peri-Alpine lakes (e.g., in 2015 in Lake Geneva; Haltiner et al., 2022). Despite co-occurrent, zebra and quagga mussels exhibit distinct invasion patterns. Zebra mussels tend to reach peak population size more quickly (2.5 ± 0.2 years) than quagga mussels (12.2 ± 1.5 years) (Karatayev et al., 2011). However, in deep lakes, quagga mussels often replace zebra mussels after 9 or more years of coexistence, attributed to their greater energetic efficiency and ability to allocate more energy to growth and reproduction (Karatayev et al., 2015; Nalepa et al., 2010; Mills et al., 1999).

To fully understand the colonization processes of quagga and zebra mussels in lakes, a detailed analysis of their reproductive dynamics is essential, including reproduction periodicity and larval dispersal. The reproductive characteristics of the zebra mussel have been globally more studied than tho*se of quagga mussel* (Karatayev & Burlakova, 2025). Dreissenid mussels reproduce essentially through sexual reproduction, exhibiting distinct sexes and very low rates of hermaphroditism (Nichols & Kollar, 1991; Spidle et al., 1995). The maturation of male and female gametes is relatively synchronous (Bachetta et al., 2010), allowing for external fertilization, with sperm being mobile in water (McMahon & Bogan, 2001). Dreissenids exhibit high fecundity, with individual females capable of releasing over 30,000 eggs during a single reproductive period (Stanczykowska, 1977) and producing more than one million oocytes per year (Walz, 1978; Borcherding, 1991). The oocyte, once fertilized in the water, transforms rapidly (2-9 days) into a veliger larva, with the development of a velum, a larval organ of feeding and locomotion (Ackerman et al., 1994). The veliger larva drifts with the water flow for at least 5 days and up to 5 weeks (reviewed in Pollux et al., 2010). Dreissenid veligers are typically concentrated in the upper layers of the water column (Sprung, 1993; Bially and MacIsaac, 2000; Wacker and Von Elert, 2003), but increased mixing or downwelling can transport them to deeper waters (Wacker and Von Elert, 2003), or result in a more uniform vertical distribution (Barnard et al., 2003). The reproductive cycle of the zebra mussel typically features two distinct peaks annually: one in spring (June to mid-July) and another in autumn (August to late October) (Claudi & Mackie, 1993). In contrast, research indicates that under optimal environmental conditions, quagga mussels can initiate reproduction earlier than zebra mussels, at greater depths and may even reproduce continuously throughout the year (Karatayev et al., 2015; Mills et al., 1996; Ram et al., 2011; Roe & MacIsaac, 1997; Claxton & Mackie, 1998; Nalepa et al., 2010).

Despite being genetically distinct (Stepien et al., 1999, 2002; Gelembiuk et al., 2006), quagga and zebra mussels are challenging to distinguish morphologically (e.g. Grigorovitch et al., 2008). DNA-based approaches, including environmental DNA (eDNA) analysis, are therefore particularly valuable for accurate identification of these species (e.g., Baldwin et al., 1996; Claxton et al., 1997; Stepien et al., 1999). eDNA methods have revolutionized biodiversity monitoring, particularly in aquatic ecosystems, and have been widely applied to detect various invasive species, including crustaceans (Geerts et al., 2018), fishes (e.g., Adrian-Kalchhauser & Burkhardt-Holm, 2016; Takahara et al., 2013), amphibians (e.g., Everts et al., 2022; Lin et al., 2019), and molluscs such as dreissenids. Numerous studies have employed eDNA from water samples to effectively detect quagga and zebra mussels (e.g. Penarrubia et al., 2016; Amberg and Merkes, 2016), offering both early detection of invasion fronts (Gingera et al., 2017) and the ability to detect low abundances (e.g., DeVentura et al., 2017; Blackman et al., 2020). DeVentura et al. (2017) also demonstrated a strong correlation between visual observations and qPCR signal quality for estimating in-place densities in the Rhine basin.

As the two species are morphologically indistinguishable during their larval stages (Grigorovitch et al., 2008), eDNA approaches are particularly relevant for characterising their reproduction in environments where they cohabit (which is often the case in colonized lakes). Here, we investigate the effectiveness of eDNA methods for monitoring the reproductive periods of the zebra and quagga mussels in lakes, which, to our knowledge, has never been explored. Sampling is a crucial step in the implementation of eDNA analyses, and for each new application it is key to ensure that sampling design is appropriate to detect the target species DNA (e.g., Ruppert et al 2019; Wilcox et al., 2018). eDNA studies in aquatic environments have largely focused on collecting and isolating DNA from water for the detection of macro-organisms eDNA (e.g. Ficetola et al., 2008; Diaz-Fergusson et al., 2014; Foote et al., 2012). However, bulk-sample approaches, that notably allows to collect large amounts of biological material, may represent a better alternative for the detection of certain taxa (e.g., Doloiras et al., 2023, Suter et al., 2020; Choi et al., 2025), including zebra mussel according to Miller et al. (2024). Therefore, we compared three eDNA sampling strategies to assess their effectiveness in detecting veliger larvae presence and abundance, and consequently, to infer the reproductive activity of dreissenid mussels in lakes. The eDNA sampling strategies included integrated water samples (0-50 m; where the veliger larvae are traditionally collected for traditional biomonitoring), sub-surface water samples, a simple and commonly used method to monitor macro-organisms’ diversity (e.g., Spear et al., 2021; Peixoto et al., 2023)), although debates exist and deeper samples may yield different biodiversity results compared to surface samples. (e.g., Turner et al 2015; Sahu et al 2025), and plankton bulk samples collected using plankton nets (0-50 m) which correspond to the sampling protocol applied to collect veliger larvae for microscopic counts.

Quagga and zebra DNA were quantified using droplet digital PCR (ddPCR), a powerful technology for detecting and quantifying rare DNA targets (e.g., Wood et al., 2019; Brys et al., 2020; Doi et al., 2015). Although eDNA approaches have been used to study dreissenid mussels (e.g., Penarrubia et al., 2016; Amberg and Merkes, 2016), ddPCR has not been previously employed for this purpose. Therefore, we developed novel primer and probe sets optimized for simultaneous multiplex use in ddPCR and validated their effectiveness and specificity to distinguish both species and to be used with eDNA samples. We hypothesize that eDNA approaches coupled with the sensitivity of ddPCR could be used to characterize the respective reproductive dynamics of quagga and zebra mussels in lakes, as it has already been demonstrated for certain fish species (Vautier et al., 2023).

The three eDNA approaches were tested in four peri-alpine lakes: Geneva and Bourget (long term colonized by zebra mussels and recently by quagga mussels), and Aiguebelette and Annecy (colonized only by zebra mussels, according to current knowledge).

## Material and methods

### Sampling sites

Sampling was conducted in four deep lakes, located in the Northern Alps—Lake Geneva (Léman), Lake Bourget, Lake Annecy, and Lake Aiguebelette—over a one-year period from August 2022 to July 2023 (12 sampling dates for lakes Bourget and Annecy; 11 for Geneva and 6 for Aiguebelette, see details below). These lakes, while geographically close, exhibit distinct ecological characteristics and varying colonization histories with respect to zebra (*Dreissena polymorpha*) and quagga (*Dreissena rostriformis bugensis*) mussels. Zebra mussels have been present in these lakes since at least the 19th century (Beisel & Lévêque, 2010), whereas quagga mussels represent a more recent invasion. Notably, quagga mussels were first observed in Lake Geneva in 2015 (with an estimated introduction around 2013) and have since become the dominant mussel species (Lods-Crozet & Chevalley, 2018; Labbat, 2020). Quagga mussels have been observed in Lake Bourget since 2019 (CISALB, personal communication). When this study was conducted, quagga mussel presence had not yet been recorded in Lakes Aiguebelette and Annecy. Within each lake, sampling locations was aligned with established long-term ecological monitoring sites, i.e., central sampling point representative of the pelagic zone monitored by OLA – Lake Observatory (Rimet et al., 2020). This sampling points are at the deepest point of each lake, historically chosen as a reference point for monitoring (Figure 1).

**Figure 1.**
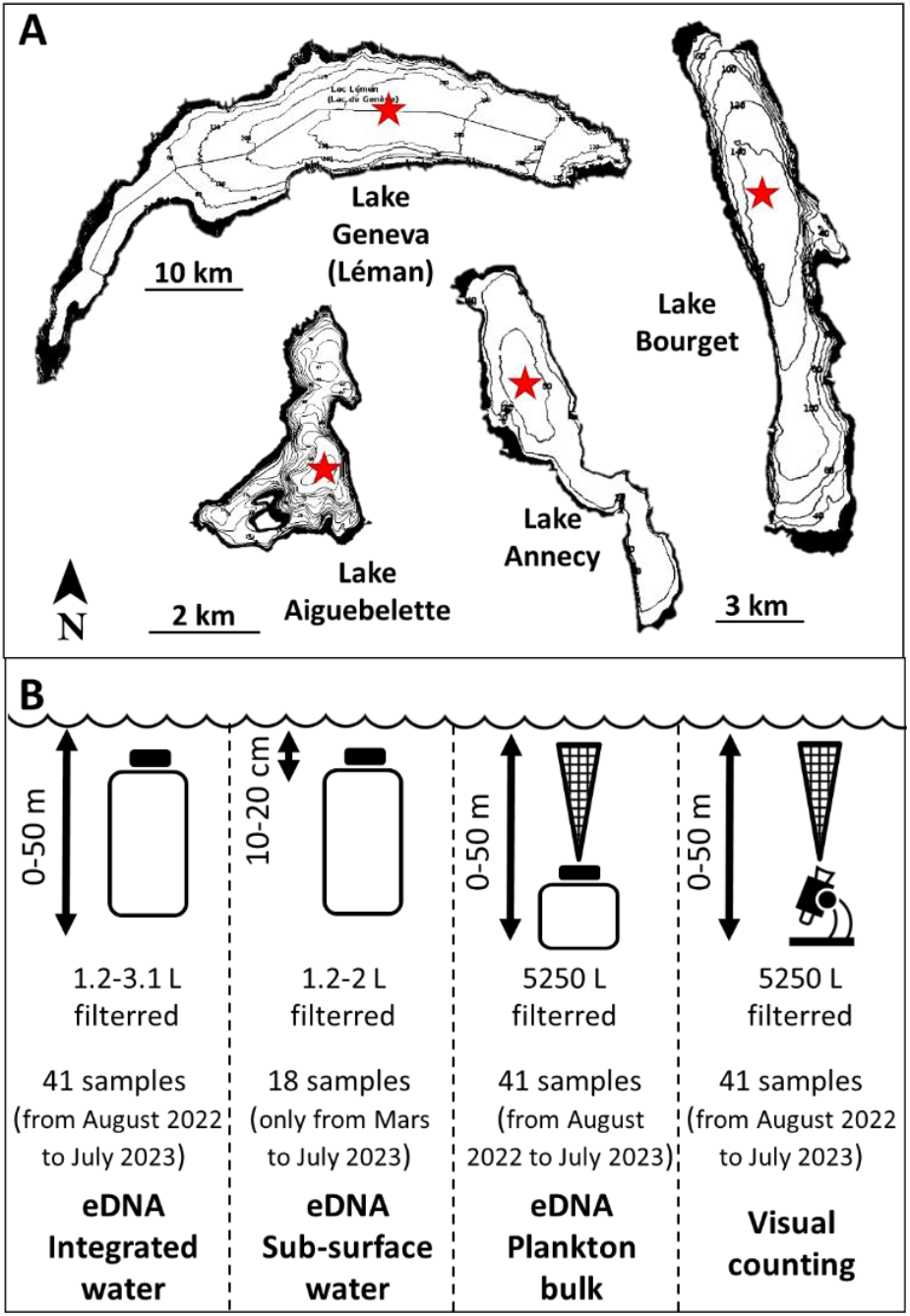
Sampling sites and strategies. **A** Maps of Lake Geneva, Lake Bourget, Lake Annecy and Lake Aiguebelette adapted from Navionics. The red stars represent the sampling sites for each lake (water and plankton nets). **B** The four types of samples collected on each lake: (i) eDNA integrated water with 1.2 to 3.1 L of water filtered per sample (41 samples) from August 2022 to July 2023, (ii) eDNA sub-surface water with 1.2 to 2 L of water filtered per sample (18 samples) from March 2023 to July 2023, (iii) eDNA plankton bulk with 5250 L of water filtered per sample (41 samples) from August 2022 to July 2023,.(ivi) Visual counting of veliger larvae from plankton bulk with 5250 L of water filtered per sample (41 samples) from August 2022 to July 2023. Lakes Bourget, Annecy and Geneva were sampled every month for one year (August 2022 to July 2023), with the exception of November for Lake Geneva due to bad weather conditions. Lake Aiguebelette was sampled only six times during the year (the usual monitoring frequency for this lake).

Lake Geneva, situated at an altitude of 372 m, has a surface area of 580.03 km^2^ and a maximum depth of 309 m. Samples were collected monthly at the deepest point of the lake (SHL2: E 6° 35’ 19.4, N 46°27’ 9.7), except in November 2022, due to unfavourable weather conditions. Lake Bourget (45°44’81 N; 5°51’36 E) located at an altitude of 372 m, has a surface area of 44.5 km^2^ and a maximum depth of 147 m. The sampling site (Point B: E 5° 51’ 35.7’’, N 45°44’ 49.7’’) is situated in the center of the lake, at its deepest point (147 m). Monthly sampling was conducted at this site. Lake Annecy (45° 51’ 7.05”N; 6° 10’ 41.92”E) located at an altitude of 447 m, has a surface area of 27 km^2^ and a maximum depth of 78.7 m. The sampling site is located in the “Grand Lac” (X=897009.793; Y=104060.45) and was sampled monthly. Finally, Lake Aiguebelette (45° 32’ 55.38”N, 5° 47’ 54.48”E) at an altitude of 374 m, has a surface area of 540 ha and a maximum depth of 70 m. The sampling site (Point A: N 45°33.040’ – E 5°48.054’) has a depth of approximately 65 meters and was monitored through six sampling campaigns (September and November 2022, and February, May, June, and July 2023).

### Sampling methods

At each sampling date, four types of samples were collected: (i) two net tows sampling were carried out, one for visual counts of veliger mussel larvae and one for the analysis of bulk eDNA on all biomass collected with net tows, and (ii) two other sampling to collect eDNA from water, one integrated sample between 0 and 50 m depth with an Integrating Water Sampler IWS III from Hydro-Bios, and one sub-surface (∼10-20 cm) sample collected by hand using a bottle (Figure 1). Sampling was carried out from August 2022 to July 2023, except for the sub-surface water samples, which only started in March 2023 (only 5 month of sampling).

### eDNA Water filtration

Water samples were stored in containers that had been previously washed in a laboratory dishwasher with an acid cycle, decontaminated with 10% hydrogen peroxide and rinsed three times with milli-Q water. The containers were kept in coolers until returning to the laboratory (max. 4h), where they were either immediately filtered, or placed in a cold room at 4°C prior to filtration. Although immediate filtration was preferred, in some cases filtration had to be postponed until the following morning due to late return from the field. The maximum conservation time at 4°C prior to filtration was approximately 14 hours. In total, 59 water samples were processed (41 integrated water samples and 18 sub-surface water samples). Filtration of the samples was performed in the laboratory using Sterivex TM MILLIPORE filtration units (0.45 µm porosity) according to the detailed protocol of Vautier (2024a). The filtration units were stored without preservation buffer, directly frozen at -80°C. Between 1.2 L and 3.1 L of water was filtered per filtration unit. Control samples, known as field controls, were also processed. These were decontaminated bottles of DNA-free water that were opened in the field during sampling, then sealed and placed in a cooler with the other samples. The control samples were then processed in the same way as the other water samples. Control samples were taken at regular intervals during the different sampling campaigns, with a total of 8 control samples (2 per lake).

### Sampling of plankton bulk

The vertical net tows were used to collect bulk samples between 0 and 50 m, using a net with a porosity of 64 µm and a diameter of 36,5 cm (the volume of water filtered was therefore 5250 L of water per net tow). Each sample collected was then stored in a 250 ml glass bottle, which was placed in a cooler until returning at the laboratory where the samples were frozen at -80°C. The volume of bulk samples varied from 178 to 245 mL. For each sampling, two vertical net tows were collected, one dedicated to microscopic observation and counting and the other to bulk-eDNA analysis. In total, 82 bulk samples were collected, 41 for microscopic counts, and 41 for eDNA analysis.

### Counting of veliger larvae

For each sample (net tow), the number of veliger larvae was counted using a binocular loupe at approximately X10 magnification and 200 µL subsamples were taken from the 250 ml bottle containing the entire net contents previously decanted (volumes from 3 mL to 15 mL). Subsamples were counted as long as no larvae was observed. If no larvae were observed, the entire bulk was counted. The entire sample volume was examined for 10 samples out of 41 bulk samples dedicated to microscopic counts. For the other samples, an average three 200 µL sub-samples were counted (ranging from 1 to 22 sub-samples, depending on larval density). Between 0 to 67 individuals (larvae) were counted in the analyzed sub-samples. The number of larvae was then expressed as number of individuals per filtered litre (considering the total volume of water filtered through the net tow i.e., 5250 litres of water).

### Extraction of eDNA from water

DNA was extracted from the filtration units using the NucleoMagDNA/RNA Water Kit from Macherey Nagel, and the MagnetaPure 32 nucleic acid purification system from Dutscher. The extraction process was conducted in accordance with the detailed protocol of Vautier et al. (2024b). The extracted DNA was eluted in 50 µL of DNA-free water, quantified with a Nanodrop spectrophotometer, and stored at a temperature of -20°C prior to ddPCR analysis. Extraction controls were also conducted, whereby the lysate utilised for the initial extraction was replaced with C1 buffer from the NucleoMagDNA/RNA Water Kit. A total of four extraction controls were produced.

### Extraction of eDNA from plankton bulk

Samples frozen at -80°C were thawed at 4°C, after which a decantation step was carried out to concentrate the biomass contained in the samples. The funnels were previously decontaminated with 10% hydrogen peroxide and rinsed three times with milli-Q water. Subsequently, the biomass was centrifuged at 5000 g for one minute to remove as much supernatant as possible. The total volume of biological material thus concentrated varied between samples, with a range of 0.3 to 5.9 ml. The extractions were performed using 100 µL of this total volume. A correction of the ddPCR results was subsequently performed to account for the fact that the total volumes of biological material from which the 100 µl aliquot was taken varied among samples (see” ddPCR analysis” section). The NucleoSpin Tissue kit from Macherey Nagel was used to perform the extractions in accordance with the manufacturer’s protocol. Pre-lysis with proteinase K was carried out for two hours. Following this, the DNA was eluted in 100 µL of BE elution buffer and quantified with a Nanodrop. The DNA was then preserved at -20°C prior to ddPCR analysis. Extraction controls were also performed, in which case the 100 µL of biomass was replaced by 100 µL of DNA-free water. A total of six extraction controls were produced.

### Development and validation of primers and probes

Primers and probes targeting *Dreissena polymorpha* and *Dreissena rostriformis bugensis* were developed and validated following the protocol detailed by Vautier et al. (2023). The DNA sequences of the COI gene for these two species have been collected from the Barcode of Life Database (Ratnasingham and Hebert, 2007). PCR primer-probe sequences targeting quagga and zebra mussels COI gene were designed using the Primers3 software (Untergasser et al., 2012). *In silico* validation was performed in order to select primer-probe sets that would theoretically amplify no other species (i.e. no perfect match in the NCBI databases) and that would exhibit the greatest number of nucleotide differences between the quagga and zebra species. The specificity of the primers selected was then tested on DNA extracted from tissues of various species of molluscs, gastropods and crustaceans present in the lakes studied (*Anodonta cygnaea, Sphaerium corneum, Ampullaceana balthica, Lymnaea stagnalis, Gammarus spp*., *Hemimysis anomala, Orconectes limosus and Pacifastacus leniusculus*), in addition to the two species of dreissenids studied in this article. The primers and probes selected are presented in Table 1. The amplicon sizes for quagga and zebra are 97 and 153 nucleotides respectively.

**Table 1.**
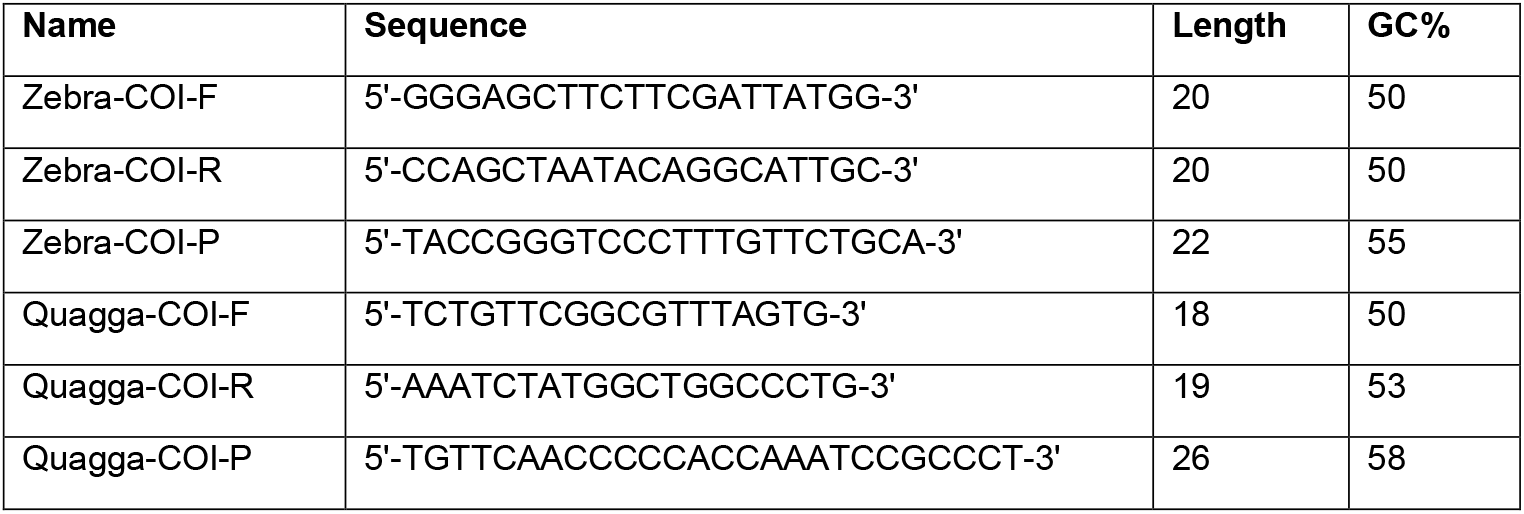
Names, sequences and characteristics of primers and probes developed. F-Forward, R-Reverse and P-Probe.

### ddPCR analysis

The ddPCRs were performed in duplex, simultaneously targeting the mitochondrial genomes of the two mussel species. The fluorescent markers chosen for the hydrolysis probes were FAM (∼517nm) for zebra mussels and HEX (∼556nm) for quagga mussels. The eDNAs extracted from each sample were analyzed independently. Negative controls were field controls, extraction controls and DNA-free water, while positive controls were DNA extracted from tissue material of each target species.

The ddPCRs were conducted using the Bio-Rad QX600 ddPCR system (Bio-Rad, Temse, Belgium) in a total volume of 20 µL, in accordance with the protocol of Vautier et al. (2024c). Each reaction contained 1x Bio-Rad ddPCR supermix for probes (without dUTP), 900 nM of each primer, 250 nM of probe, between 4 µL of template DNA, 5 U of AflII restriction enzyme and was topped up with diethylpyrocarbonate water (DEPC) (Sigma-Aldrich, Overijse, Belgium). Twenty microliters of the DNA/reaction medium mixture were transferred to the sample compartments of the Droplet Generator DG8 cartridges (Bio-Rad, Cat. No. 1864008), and 70 µL of Droplet Generation Oil for Probes (Bio-Rad, Cat. No. 186-4005) was added to the appropriate wells.

Subsequently, the droplets were analysed using a QX600 droplet reader (Bio-Rad). All droplets were subjected to fluorescence analysis using QX Manager Software 2.0. The fluorescence amplitude threshold, employed to differentiate between positive and negative droplets, was established manually by the analyst as the midpoint between the mean fluorescence amplitude of the positive and negative droplet groups. The same threshold was applied to all wells of a given PCR plate. The mean number of accepted droplets was approximately 17,000.

To estimate eDNA concentrations, expressed as copy numbers per liter of filtered water, the formula proposed by Vautier et al. (2023) was employed. For eDNA samples obtained from bulk, the concentration calculation was also corrected by the total volume of the bulk sample after decantation and centrifugation. This volume was variable between samples (from 0.3 to 5.9 mL), but the quantity of the bulk sample collected for DNA extractions was fixed (100 µL).

### Statistical analysis

Spearman correlations and t-tests were performed using the R statistical computing environment (version 4.3.2).

## Results

### Primer and probe sensitivity and specificity tests

The LOD and LOQ were estimated from dilutions of DNA extracted from tissues for both newly designed primers set. The LOQ for both primers set was obtained at 0.4 pg of DNA, for respectively a mean of 1.49 copies per reaction for quagga and 3.21 copies per reaction for zebra. The LODs were obtained at different DNA quantities, for quagga at 0.08 pg DNA for a mean of 0.30 copies per reaction, and for zebra at 0.016 pg of DNA for a mean of 0.19 copies per reaction (Figure 2).

**Figure 2.**
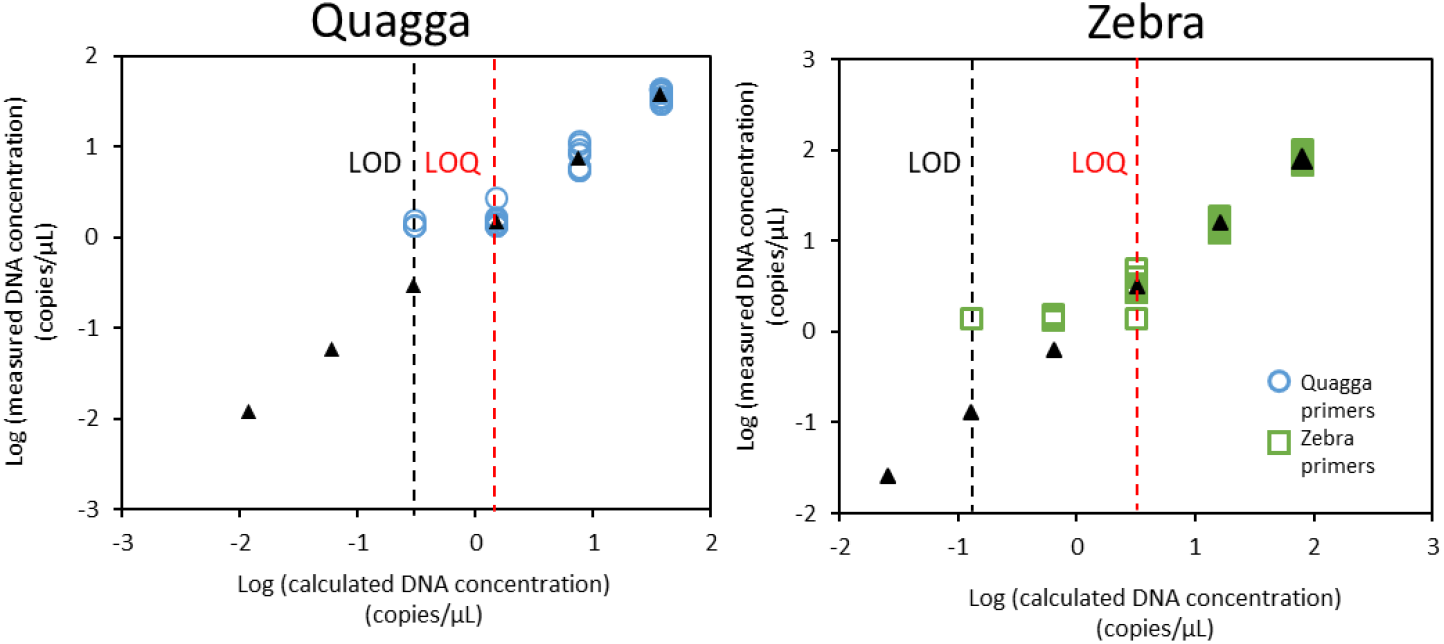
Sensitivity, limit of detection (LOD) and limit of quantification (LOQ) of primer sets. The sensitivity and limits of detection and quantification of the primer sets were evaluated using six amounts of DNA (10, 2, 0.4, 0.08, 0.016 and 0.0032 pg), obtained from five successive dilutions of the highest amount of DNA (10 pg) for quagga and zebra mussels with 10 replicates each. Black triangle = theoretical estimate of the number of target gene copies per ddPCR well, based on the number of gene copies obtained from the highest amount of DNA. Quagga primers (blue circle), zebra primers (green square).

Primer specificity was determined by dPCR with 10 pg of DNA extracted from tissues of 8 other species (molluscs, gastropods and crustaceans) present in the studied lakes, in addition to quagga and zebra mussels. No aspecific amplification was observed, leading to the conclusion that the newly designed primers are specific to each mussel targeted.

### Monthly dynamics of eDNA concentration and number of veliger larvae in the four lakes

The presence of veliger larvae was revealed through visual counts in all four lakes. In two of these lakes, Geneva and Bourget, the presence of larvae was observed throughout the year, while no larvae were observed in lakes Annecy and Aiguebelette between January and May. The highest number of larvae was observed during the ‘warm’ period, from June to October, in all four lakes.

With regard to the eDNA analysis, both quagga and zebra mussels were identified at least once in each of the four lakes (Figure 3). A quagga signal was detected in all samples from Lakes Bourget and Geneva, while it was only present in 7 out of 29 samples from Lake Annecy and 5 out of 15 from Lake Aiguebelette. zebra was successfully identified in only 2 out of 27 samples from Lake Geneva, and this was only with bulk eDNA samples. For Lake Bourget, 12 out of 29 samples exhibited the presence of zebra, predominantly during the warm period for all eDNA approaches. 14 out of 29 samples were positive for zebra in Annecy and 4 out of 15 in Aiguebelette; in these two lakes, the positive signals for the two mussel species occurred predominantly during the warm period.

**Figure 3.**
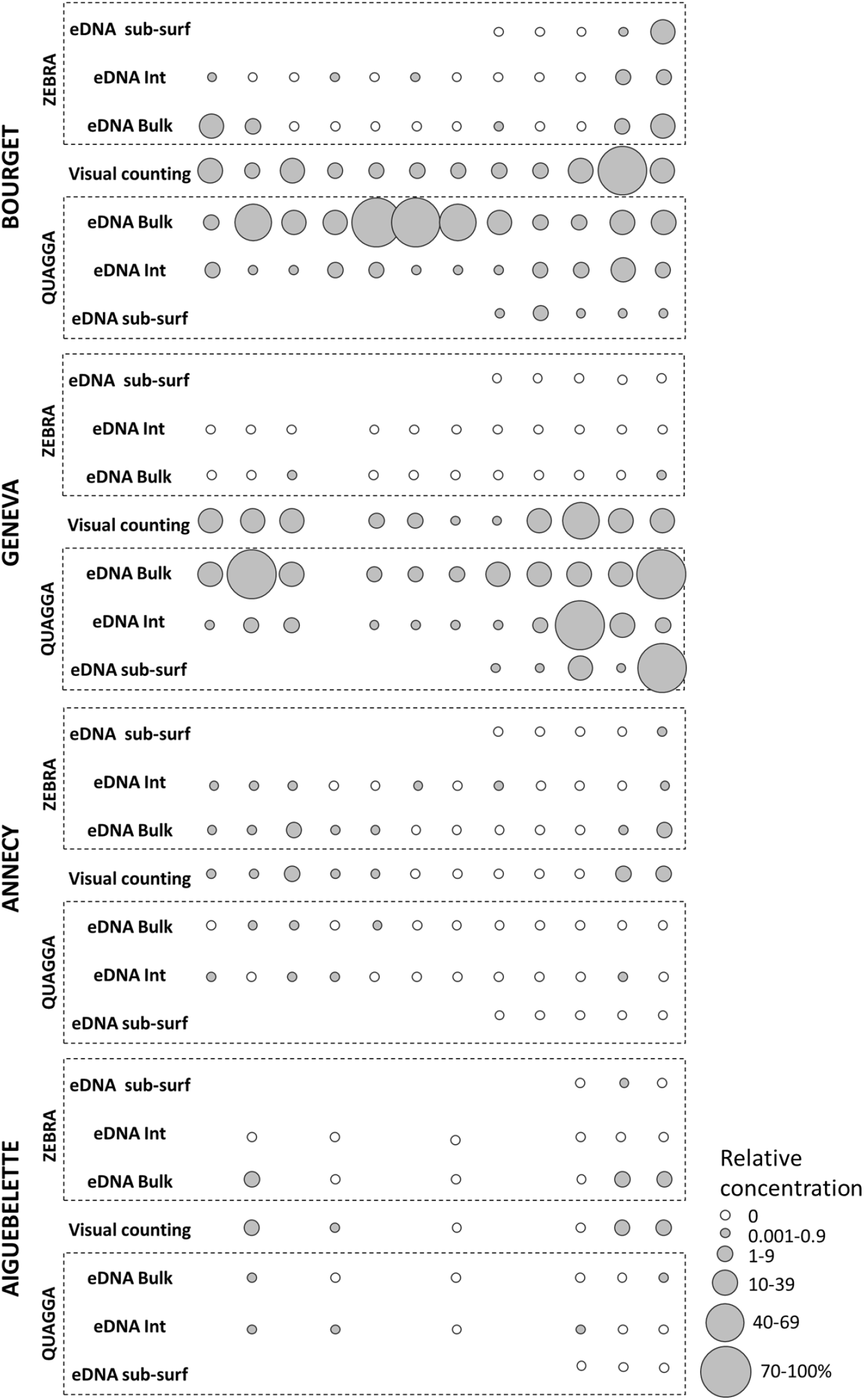
Monthly dynamics of eDNA concentrations and number of veliger larvae over one year in the four studied lakes (Geneva, Bourget, Annecy and Aiguebelette). The three eDNA approaches are labelled as follows: water integrated between 0 and 50 m depth (eDNA Int), sub-surface water (eDNA sub-surf) and plankton bulk (eDNA Bulk), and are presented for the two species quagga and zebra. The microscopic counts of veliger larvae in plankton bulk (visual counting) are undifferentiated for the two species. The results are expressed as relative eDNA concentrations or relative numbers of larvae for the counting, using as a reference (100%) the highest concentration of eDNA, or the highest number of larvae counted. If there is no circle, no sample has been taken.

For Lake Geneva, for all eDNA approaches, the quagga and zebra eDNA signals were stronger during the warmer period, while for Lake Bourget, a similar trend was found only for water-based eDNA approaches, and a stronger signal was obtained for the quagga with the bulk-based approach during the colder period.

A comparison of the ddPCR results of the 82 integrated water eDNA samples with the 82 bulk DNA samples reveals that 61 exhibit identical results in terms of presence/absence, i.e. 74% of cases. For 12 ddPCR results, bulk eDNA detected the target but integrated water eDNA did not, and for 9 the inverse was true. The detection efficiency of both approaches was found to be comparable, with observed variations occurring only under rare signals (always below a threshold of 10% of the maximum values measured by type of sample). This comparison was not made with sub-surface water due to the fact that samples were only collected over a period of five months for sub-surface water eDNA.

False positive results were observed with integrated water eDNA (eDNA signal measured when no larvae were observed), while no such cases were observed with bulk eDNA. This suggests that bulk eDNA is more representative of the presence of larvae than integrated water eDNA. When examining the detection percentage (i.e. percentage of samples positive for eDNA detection of one of the two mussels, or, for the observation of at least one larva relative to the total number of samples), it appears that integrated water eDNA, larval counts, and bulk eDNA yield very similar results, with 82.9%, 85.5%, and 80.5% respectively. The value obtained for sub-surface water eDNA, is lower, with only 66.7% detection of one of the two mussels across all samples. Considering each species separately, it is obvious that sub-surface water eDNA consistently showed lower detection rates compared to the other two eDNA methods. The comparison of the other two eDNA methods showed that integrated water eDNA has a better detection rate for quagga mussels, with 73.2% compared to 68.3% for the bulk eDNA, while the opposite is true for zebra mussels (bulk-eDNA detects more signal with 43.9% compared to 26.8% for integrated water).

### Correlations between eDNA approaches and veliger larvae microscopic counts

As counts based on morphological criteria cannot distinguish between the two bivalve species, we compared these values with the cumulative eDNA concentrations of the two species. Strong positive and significant correlations were found between all methods with a p-value <0.001 (Figure 4). Looking at these correlations for the two mussel species separately, it appears that there is also a significant positive correlation between all methods for quagga with a p-value<0.001, except between visual counts and sub-surface water eDNA where the p-value<0.01. For zebra, the correlation coefficients are much weaker, with no significant positive correlation between eDNA and visual counts. Significant positive correlations were still found between the eDNA approaches, but with lower coefficients and p-values than those obtained for quagga. The presence of veliger larvae in the 4 lakes studied is therefore more strongly correlated with the quagga eDNA signal than with zebra mussel eDNA signal.

**Figure 4.**
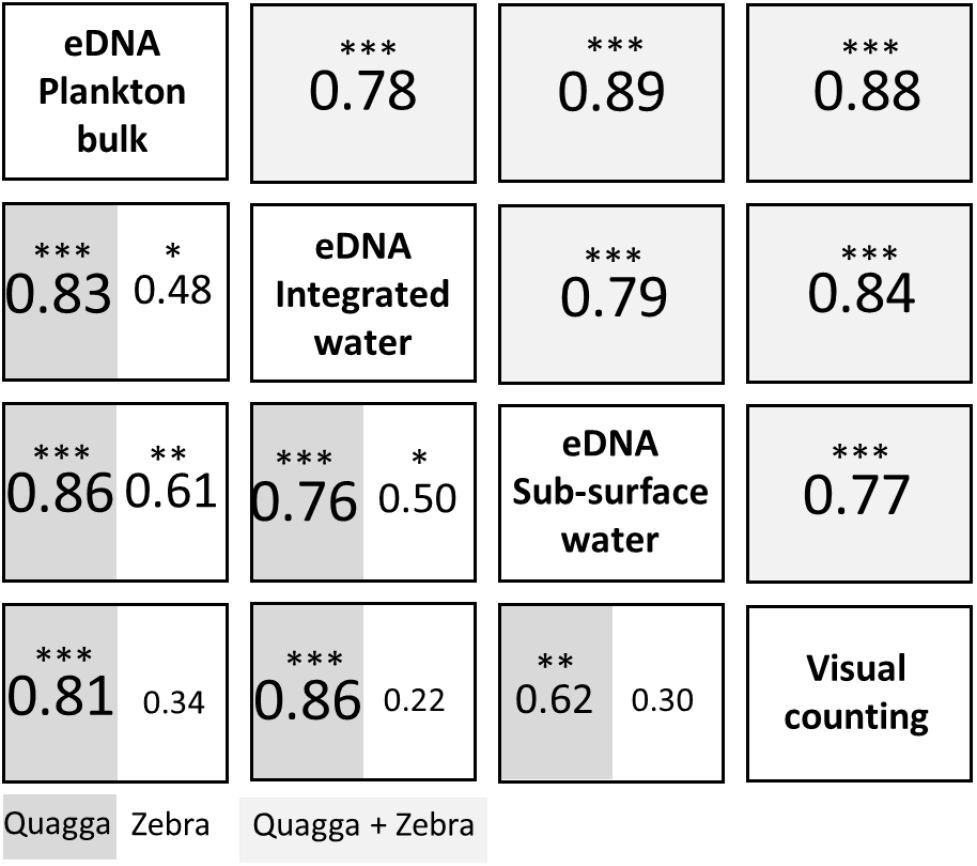
Correlations between eDNA approaches and visual counts of veliger larvae. Spearman coefficient correlations between morphological counts and eDNA concentrations obtained for zebra and quagga mussels combined (light grey top right) or separated (dark grey and white lower left) and for the four lakes combined, Geneva, Bourget, Annecy and Aiguebelette. ***: p.value<0.001, **: p.value<0.01, *: p.value< 0.05.

### Comparison of eDNA concentrations obtained for the two species in the four lakes

A comparison of the eDNA concentrations obtained with the three different eDNA approaches reveals that for the quantification of quagga, the bulk eDNA and integrated water eDNA approaches are similar, but that the sub-surface water eDNA approach yields overall lower concentrations (Figure 5). This difference is statistically significant for Lake Bourget. For zebra eDNA concentrations, no significant difference was found between the three different eDNA approaches, although the bulk eDNA approach appeared to give higher concentrations overall. This approach was also the only one to detect the zebra’s eDNA signal in Lake Geneva and in Lake Aiguebelette while the integrated and sub-surface water eDNA approaches did not (except for sub-surface water eDNA approaches in Lake Aiguebelette).

**Figure 5.**
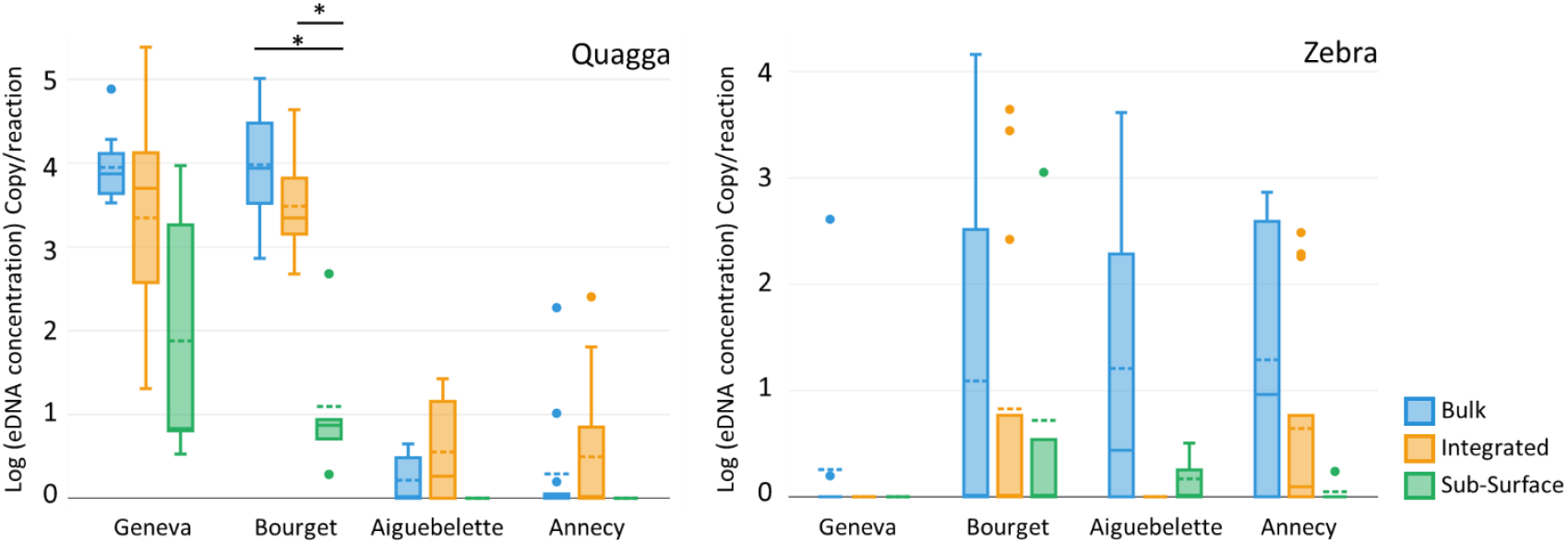
eDNA concentrations obtained for the four lakes and with the three eDNA approaches. eDNA concentrations are expressed as copy numbers per ddPCR reaction with log transformation. The bulk eDNA approach is shown in blue, integrated water eDNA in orange and sub-surface water eDNA in green. Student’s t-tests were performed between eDNA approaches within the same lake and significant differences with a p-value<0.001 are indicated with *.

## Discussion

This study provides significant advancements in both technical approaches for detecting invasive dreissenid mussels and our understanding of their ecology in peri-alpine lakes.

### Technical advances for dreissenids monitoring in lakes

This study proposes two significant technical advances for the identification and the monitoring of invasive dreissenid mussels using eDNA. Firstly, we developed new primers and probes specifically designed for multiplex ddPCR to discriminate zebra and quagga mussels; a critical advancement for monitoring these invasive species, particularly at larval stages when morphological discrimination is impossible. These primers demonstrated high specificity and sensitivity, with calculation of there LOD and LOQ enabling robust analysis of the results obtained by ddPCR. The use of ddPCR in our study represents an improvement in comparison to traditional qPCR methods which have already been shown to be effective in detecting dreissenids (e.g. Gingera et al., 2017, DeVentura et al., 2017, Blackman et al., 2020). ddPCR offers advantages such as absolute quantification without the need for standard curves (Vogelstein & Kinzler, 1999), a reduced detection limit, higher sensitivity, accuracy and reproducibility (Hindson et al., 2013) and greater tolerance to inhibitors (Hoshino & Inagaki, 2012). Furthermore, ddPCR has been shown to effectively detect rare DNA in a background of high numbers of non-target nucleic acids (Pohl & Shih, 2004), thereby enhancing the sensitivity of eDNA approaches for rare aquatic organisms’ detection (Doi et al., 2015). These benefits make ddPCR a promising tool for future invasive species monitoring programs, allowing an early detection, and then increasing the chances of successful eradication or containment, and reducing costs, owing to the fact that there is no requirement for standard curves or replicates.

Secondly, we compared three different eDNA sampling methods to monitor quagga and zebra reproduction periods in lakes: integrated water samples, sub-surface water samples, and plankton bulk samples. Peaks of eDNA, likely resulting from sperm release and spawning aggregations, have been used to determine the timing of fish spawning (Vautier et al., 2023; Bylemans et al., 2017; Tsuji & Shibata, 2021; Takeuchi et al., 2019). Such eDNA spikes signify that this life history stage is particularly fruitful for monitoring by eDNA. In this study, we hypothesised that analogous eDNA peaks would be observed during the reproductive cycle of mussels, and mainly attributable to the presence of veliger larvae into the water. Our results demonstrate that the three eDNA sampling methods are effective in estimating mussel reproductive periods in lakes. This study is, to the best of our knowledge, the first to demonstrate the applicability of eDNA to monitor mussel reproduction, with the advantage of distinguishing between quagga and zebra mussels spawning signal. All three eDNA sampling approaches showed significant positive correlations with veliger counts conducted in the four studied peri-alpine lakes. However, notable differences were observed among the methods:

Sub-surface water eDNA consistently proved less effective than the other two eDNA approaches in detecting the presence of either mussel species both in terms of % of detection and in terms of quantity (i.e., weaker signal with sub-surface sampling). This low efficiency is particularly noticeable for weak eDNA signals, i.e. sub-surface water eDNA failed to detect quagga mussels in Lakes Annecy and Aiguebelette whereas the other two eDNA approaches identified a weak signal. These results are consistent with those reported by Miller et al. (2024), who demonstrated that the use of eDNA bulk samples is more effective for detecting the eDNA signal from zebra in lakes than sub-surface water eDNA samples. While this method (sub-surface sampling) is the simplest to implement, it is the least effective for quantifying dreissenid reproduction signals in lakes using the protocol applied here. Methodological optimizations, such as increasing the filtered water volume, could potentially enhance the performance of our approach for sub-surface sampling. In this study, we filtered a maximum of only 2 L per sample, previous research has shown that filtering larger volumes of water can lead to greater biodiversity detection for certain taxa, such as metazoans (e.g., Govindarajan et al., 2022; Peres & Bracken‐Grissom, 2025).

The integrated water eDNA and bulk eDNA both demonstrated significant positive correlation with larval counts and yielded high mussels eDNA concentrations. This corroborates one of our initial hypotheses that the eDNA peaks occurring during mussel reproduction due to the release of larvae into the water, should be efficiently detectable either in bulk eDNA samples, which directly capture larvae if the mesh size (i.e. size of veliger larvae between 70 and 300µm), or in integrated water eDNA samples taken directly in the dispersal area where the larvae are present. However, while bulk-based eDNA approach is robust for qualitative presence/absence veliger larvae information, with a positive eDNA signal only when larvae are observed, two false positive results were observed with integrated water eDNA (eDNA signal when no larvae were observed). This is not unexpected, since eDNA extracted from water samples is derived from larvae as well as adult individuals present throughout the year (DNA signals that are free in the water column or bound to particles). A positive signal from bulk-based eDNA almost certainly indicates the presence of larvae and therefore reproduction in the previous weeks reviewed in Pollux et al., 2010), whereas this is not necessarily the case for a positive signal from water-based eDNA.

Although both approaches show a positive and significant correlation with visual counts of larvae in the four lakes taken all together, this is not the case for bulk eDNA in Lake Bourget alone (Sup. Mat. 1). Indeed, bulk-based eDNA samples showed the strongest quagga mussel signal during colder months (December, January, and February) in Lake Bourget, contradicting other approaches, including visual counts that indicated weaker signals during this period. This discrepancy is likely due to an inherent bias in bulk samples collected from plankton nets. The concentration of veliger larvae in these samples varies with the total plankton quantity in the environment, with higher concentrations in winter due to lower overall plankton quantity. When eDNA is extracted from an aliquot of this planktonic bulk, the probability of including a veliger larva in the aliquot increases when total plankton biomass is low, potentially leading to an overestimation of the larval signal in winter (bulk up to 20 times more important in summer than in winter). To mitigate this effect, eDNA data were adjusted using total bulk plankton volume. However, potential additional improvements to address this issue could include an optimization of the DNA extraction protocols to be able to extract DNA from the entire net tow biomass rather than just an aliquot, or the lysis of the entire bulk sample before taking an aliquot to homogenise the DNA. In Lake Bourget, the integrated water eDNA approach showed a positive significant correlation with visual counts and appeared to capture the reproductive dynamics more effectively than bulk eDNA.

An unexpected result is that the use of bulk eDNA provided superior detection of zebra mussels (the only method to detect them in Lake Geneva), while integrated water eDNA provided enhanced detection of quagga (detected in a greater number of samples than bulk eDNA). One factor to consider here is the fact that plankton nets, despite filtering more than 5000 L of water - which understandably gives a higher probability of capturing larvae compared to the 5 L of water collected for the integrated water samples - do not retain DNA signals that are free in the water column or bound to particles smaller than the mesh size (64 µm in this case). Numerous studies have demonstrated that the porosity employed is a pivotal factor in enhancing the likelihood of detecting eDNA signals within aquatic environments (e.g.; Kumar et al., 2022; Jo et al., 2020). This may therefore explain the lower efficiency in capturing part of the eDNA signal when a large proportion of the target eDNA signal is “found” in the < 64µm fraction. If this explanatory factor was to be considered, our results would suggest that the <64 µm eDNA fraction (the eDNA fraction not associated with veliger larvae) is proportionally more important for quagga mussels than for zebra. This finding is consistent with the sampling locations selected in this study, which were in pelagic zones and therefore closer to the typical habitat of quagga mussels. Quagga mussels have been observed to develop and reproduce at greater depths than zebra mussels (Roe and MacIsaac, 1997; Claxton and Mackie, 1998; Nalepa et al., 2010). It is reasonable to hypothesise that the detection of the eDNA released by the quagga (“free” DNA) will be enhanced if the sampling is conducted above their habitat in the central region of the lake. In contrast, zebra mussels have been found to be prevalent in littoral areas, and the DNA they release (with the exception of that present within their larvae) may be in lower abundance at the sampling site in the centre of the lake (further from the emission zone). Moreover, hard-shelled or shell-encased organisms are known to shed low amounts of eDNA compared to other aquatic taxa (e.g.; Adams et al., 2019; Andruszkiewicz et al., 2021; Sansom & Sassoubre, 2017); and the degradation rates of the released eDNA are considered similar to those reported in the literature for other taxa (Ruiz‐Ramos et al., 2024). There is therefore little chance of the eDNA of these species dispersing and remaining detectable far from the emission zone. Conversely, bulk eDNA has the capacity to effectively detects the littoral zebra mussel signal even when the sampling is performed at the centre of the lakes, as it could primarily originate from veliger larvae, representing a signal capable of propagating over long distances within water (Stoeckel et al., 2004; Orlova et al., 2004).

With the protocols tested in this study, the bulk-based eDNA method appears to provide superior qualitative information regarding the presence or absence of mussel larvae, and was the only method capable of detecting zebra in Lake Geneva. However, the method based on integrated water eDNA provides a very reliable estimate of the dynamics of mussel reproduction in lakes, sometimes better than with bulk eDNA, but with the presence of false positives. In light of these findings, the bulk eDNA approach is recommended when the objective is to ascertain the presence of veliger dreissenid larvae with certainty. Nevertheless, improvements to the methodology may be implemented in order to enhance consistency with reproduction dynamics. The eDNA integrated water approach has been demonstrated to be an effective alternative for monitoring the reproductive dynamics of quagga and zebra mussels in lakes; however, the possibility of false positives must be considered.

### Advancements in knowledge about the ecology of dresseinids in lakes

This study provides the first indication of quagga mussel presence in Lakes Annecy and Aiguebelette. The efficacy of eDNA in the early detection of dreissenids in lakes and rivers has been demonstrated in previous studies (e.g. Gingera et al., 2017, DeVentura et al., 2017, Blackman et al., 2020). Early detection played a pivotal role in the successful eradication of Black striped invasive mussel (*Mytilopsis sallei*) population in Australia (Bax et al., 2002), underscoring the significance of developing and implementing effective tools for the early detection of invasive species. Here we reported the first detection of quagga mussels in lakes where only zebra mussels were known to be present. The presence of the quagga has been demonstrated to have significant consequences on lacustrine ecosystems, including an increase in water clarity through the filtration action of mussels (Budd et al., 2001), alterations in phytoplankton composition (Vanderploeg et al., 2010), and damage to infrastructure (Connelly et al., 2007). It is therefore essential to establish specific management plans following the early detection of quagga in Lake Annecy and Aiguebelette, as with the prevention of the arrival of other populations, which, through the process of hybridisation, could give rise to offspring that are more vigorous or better adapted to the colonised environment (Roman & Darling, 2007; Marescaux et al., 2016). Furthermore, ongoing monitoring of these species in areas that have not yet been invaded is required.

The study also revealed interesting patterns in the reproductive dynamics of these two invasive mussels. In Lakes Geneva and Bourget, where quagga mussels have become dominant, we observed year-round presence of veliger larvae, and eDNA quantifications suggest that these larvae are quagga larvae. This contrasts with Lakes Annecy and Aiguebelette, dominated by zebra, where no larvae were detected between January and May. As dreissenids larvae remain in the water for a maximum of five weeks before settling (Sprung, 1989), our results indicate that quagga mussels could reproduce throughout the year in peri-alpine lakes, while zebra mussels appear to be limited to warmer months. This is consistent with previous research suggesting that quagga mussels are more tolerant to cold temperatures, being able to develop at 9°C (Claxton & Mackie, 1998) and even reproduce at temperatures as low as 4.8°C (Roe & MacIsaac, 1997), whereas for the zebra the limit temperature appears to be much higher, around 12°C (Sprung, 1995; Claxton & Mackie, 1998). The extended reproductive period of quagga mussels, in addition to its low winter mortality (D’Hont et al., 2018), may contribute to their competitive advantage over zebra mussels in peri-alpine lakes.

For Lakes Annecy and Aiguebelette, larvae were only observed from June to December, which may correspond more closely to the warmer breeding periods for the zebra, and these results are corroborated by eDNA, and in particular by the bulk-based approach. For the four lakes, more larvae were observed and a higher eDNA signal measured during the ‘warm’ period, from April to October, suggesting that this is the preferred breeding period for both species, as has already been shown (Claudi & Mackie, 1993).

The strong correlation between Quagga mussel eDNA concentrations and veliger larvae counts, compared to the weaker correlation for zebra mussels, further supports the dominance of quagga mussels in Lakes Geneva and Bourget. This shift in species composition has important implications for ecosystem management and underscores the need for species-specific monitoring approaches. As quagga almost completely supplants zebra in deep lakes after around 9 years (Karatayev et al., 2015, Nalepa et al., 2010), ADNe approaches are consistent with the fact that quagga arrived in Lake Geneva before it arrived in Lake Bourget (first observed in 2015 (Haltiner et al., 2022) and 2019 (Cisalb communication) respectively). The eDNA approaches therefore seem to provide a reliable representation of the dynamics between the two species and further monitoring would be interesting in order to determine whether quagga densities stabilise after around 12 years as presented by Karatayev et al. (2011).

### Perspectives

As dreissenid mussels continue to spread and impact freshwater ecosystems, the tools and knowledge gained from this research will be invaluable in developing effective monitoring and potentially supporting management plans for both invaded and non-invaded lakes.

To take these developments a step further, future research should focus on optimizing eDNA sampling and analysis methods to address current limitations in detection and quantification. In this study, eDNA sampling was carried out in the deepest zone of the lakes, and water-based eDNA showed poorer detectability of zebra compared with quagga. As zebra mussels are mainly found in shallower areas than quagga, sampling in more littoral zones could favor the capture of zebra free eDNA, and therefore its detectability. With regard to the bulk-based eDNA approach, a possible bias was observed due to the fact that the total bulk volume varies with the seasons. Extractions are carried out from a sub-sample with a fixed volume, thus increasing the probability of collecting a veliger larva in winter, and decreasing it in summer. To counterbalance this bias, optimization of extraction protocols could be achieved to extract DNA from the entire bulk, or lysis of the entire bulk to be extracted, so that the target DNA would be more homogeneously distributed in the sample when the sub-sample would be collected for extraction. The ddPCR approach developed in this study could also be adapted for monitoring other invasive aquatic species, potentially improving early detection and characterization of their reproductive cycles; therefore helping to enhance our understanding and management of invasive species.

## Acknowledgments

The authors wish to thank the OLA services – ANAEE-France for their assistance with field sampling, and in particular Pascal Perney, Jean-Christophe Hustache, and Philippe Quetin. We are grateful to Leslie Laine for performing the larval counts under the microscope. We also acknowledge Jean-Nicolas Beisel for the CIPEL-funded project on the comparison of quantification methods for quagga mussels in Lake Geneva, especially for the bibliographic synthesis he conducted. Finally, we thank Isabel Blasco-Costa from the Natural History Museum of Geneva for providing invertebrate samples used for primer specificity testing.

