## Supplemental Material 1 for "Advancing eDNA methods for monitoring the reproduction of quagga and zebra mussels in lakes"

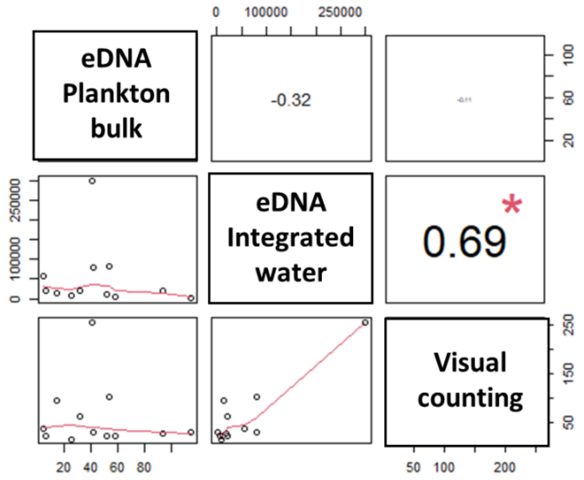


**Sup. Mat. 1.** Correlations between eDNA approaches and visual counts of veliger larvae in Lake Bourget. Spearman coefficient correlations between morphological counts and eDNA concentrations obtained for Zebra and Quagga mussels combined. ***: p.value<0.001, **: p.value<0.01, : *p.value< 0.05.
